# A neuronal theory of sequential economic choice

**DOI:** 10.1101/221135

**Authors:** Benjamin Y. Hayden, Rubén Moreno-Bote

## Abstract

Principles derived from recent studies have begun to converge and point to a consensus for the neural basis of economic choice. These principles include the idea that evaluation is limited to the option within the focus of attention and that we accept or reject that option relative to the entire set of alternatives. Rejection leads attention to a new option, although it can later switch back to a previously rejected one. The referent of a value-coding neuron is dynamically determined by attention and not stably by labeled lines. Comparison results not from explicit competition between discrete representations, but from value-dependent changes in responsiveness. Consequently, comparison can occur within a single pool of neurons rather than by competition between two or more neuronal populations. Comparison may nonetheless occur at multiple levels (including premotor levels) simultaneously through a distributed consensus. This framework suggests a solution to a set of otherwise unresolved neuronal binding problems that result from the need to link options to values, comparisons to actions, and choices to outcomes.

## INTRODUCTION

Economic choice, the selection of options based on their value, is a core process in the repertoire of intelligent organisms (Pearson et al., 2014; Rushworth et al., 2011; Rangel and Hare, 2010). Neuroeconomic research has successfully identified some of the major brain regions associated with valuation and choice, especially the orbitofrontal cortex (OFC), ventromedial prefrontal cortex (vmPFC), dorsal anterior cingulate cortex (dACC), and ventral striatum (VS). These regions are activated by the values of offers and outcomes and show correlations with comparison-related processes (Wallis, 2007, Bartra et al., 2013, Haber and Behrens, 2014; Rushworth et al., 2011). Lesion studies support the idea that these regions have a direct causative role in choice (e.g. Noonan et al., 2010, Kennerley et al., 2006; Camille et al., 2011). All measures point to some degree of specialization within these regions, although their respective roles remain debated (Rushworth et al., 2011). While the locations of value-related processing are now established, the mechanisms of choice are not. Nonetheless, we believe that a series of recent studies have begun to limn something of a consensus view – if not a model, at least a framework for one.

A common proposal is that value comparison is implemented by direct competition, via mutual inhibition, between discrete sets of neurons whose responses correspond to the value of particular options (e.g. Rustichini and Padoa-Schioppa, 2015; Chau et al., 2014; Hunt et al., 2015; Louie et al., 2011; Hunt et al., 2012; Soltani et al., 2006; Padoa-Scioppa, 2011). From this perspective, value representations are aligned to neuron identity by a *labeled line code:* a neuron’s firing rate indicates a value and its identity (its notional label) indicates which option has that value. This stable relationship makes implementing choice straightforward: the two populations compete for control of a third set of neurons and whichever set of neurons wins the competition determines the chosen option. However, this approach introduces several problems. First, it necessitates redundant reduplication of circuitry for computing value for each offer. Second it requires precise wiring to implement it (or else a well-informed supervisory system that dynamically creates that wiring.) Third, it does not readily scale up to more than two offers, nor does it deal well with newly introduced or novel offers. Fourth, it introduces the need to coordinate a flexible linkage between offers, values, actions, and positions in space; we call these problems the neuroeconomic binding problems. While these problems are undoubtedly surmountable, we wondered instead whether an alternative approach could provide a better framework for models of economic choice.

Several recent findings have, in our view, begun to bring into focus an alternative picture of how choice works. Here, we first review that evidence, with a focus on primate single unit recoding studies. First, we describe six major research trends that, together, point towards our integrated framework. We then describe a framework that is consistent with, and supported by, these research trends. This framework is also directly motivated by principles of foraging theory that, in our view, constructively interact with the principles of neuroeconomics to guide our understanding of reward-based choice (Pearson et al., 2014; Hayden, 2017). Finally, we provide a speculative discussion of the ways that our framework points towards novel solutions to old problems, especially the neuroeconomic binding problems.

## PART I: Empirical findings

### 1. We evaluate only one option at a time

Attention has a narrow bandwidth. While the spotlight of attention can in principle be split, it is difficult to do so; it is much easier to simply attend items serially (Treisman and Gelade, 1980; Egeth and Yantis, 1997). Some features of the visual scene, such as color, can be analyzed in parallel, but serial processing is greatly preferred for complex feature extraction. Value, which is typically determined by the combination of multiple features, seems likely to be the type of complex feature that requires unitary focal attention.

It is not surprising, then, that when two options are presented in the visual field, our eyes naturally shift back and forth between them to evaluate them (Krajbich et al., 2010; Orquin and Mueller Loose, 2013). When gaze is held fixed by the experimenter the spotlight of attention may nonetheless covertly shift between the two options. And when they are not presented visually, there may still be a shifting covert mental focus of attention that selects one at a time for processing. The limitation of evaluation to a single option is also consistent with ideas developed by foraging theory (Stephens and Krebs, 1986; Kacelnik et al., 2011; Kolling et al, 2012; Shapiro et al., 2008; Hayden, 2017). Specifically, foraging theorists generally point out that evolution has shaped the development of a system that will accept or reject a single option, and that when faced with multiple options, we likely perform multiple more-or-less independent accept-reject decisions (Kacelnik et al., 2011).

This gaze-dependent fixational model is supported by studies of the relationship between fixation patterns and choices (Krajbich et al, 2010; Krajbich et al., 2011). Neural evidence supports, or is consistent with, the idea that the core value regions, vmPFC, OFC, and VS encode the value of the attended option (Lim et al., 2011; Strait et al., 2014; Strait et al., 2015; Blanchard et al., 15; Xie et al., 2017; Rudebeck et al., 2013; McGinty et al., 2016). In a recent study of the OFC, ensembles of neurons alternated between encoding only one of the two available options rather than encoding both at the same time (Rich and Wallis, 2016). Notably, these coding states did not track the locus of gaze, but presumably tracked the focus of attention, suggesting that it is attention, not gaze direction *per se*, that determines which option is evaluated. The underlying foraging models that support these ideas have also proven quite useful in explaining a great deal of brain activity (Kolling et al., 2012; Kolling et al., 2014; Hayden, 2017; Blanchard and Hayden, 2014; Boorman et al., 2011, Boorman, et al., 2013, Boorman et al., 2009).

*Implications for the framework:* If we can only attend one offer at a time, then processing of the two offers in a binary choice must occur serially, not in parallel **(Figure 2)**. (The same is true for choices with more than two offers, see below). Relative to parallel models, serial processing poses a new problem and solves an old one. The new problem is that it requires a working memory buffer so that the value of a previously attended option can be maintained in order for any comparison to occur. The solved problem is the option-value binding problem. Because attention is limited to one option, there is no ambiguity about the reference of value-related neural responses. As long as the decoder knows where the focus of attention is, the referent of the value signal is unambiguous (that focus need not be spatial; it may be abstract and conceptual.)

### 2. We decide whether to accept or reject that option

If only one option is attended at a time, it is natural that the decision will be simply to accept or reject that one. Rejection would be favored, even for very good options, when the cost of inspecting the next one is low and there is no cost to returning to the first one (as in most laboratory binary choice tasks, although not necessarily as in natural contexts). In the laboratory, then, we would therefore expect a period of inspection before a period of choice. As noted, foraging theory has long emphasized the idea that preys are naturally encountered alone, and thus our brain’s evolved choice strategy is to either accept or reject a single offer (Stephens and Krebs, 1986; Charnov, 1976; Krebs et al., 1977). This decision is made relative to an estimate of the value of rejection (i.e. the opportunity cost of accepting), known as the *background value*.

From this accept-reject perspective, ostensibly binary choices involve two largely distinct accept-reject decisions, one for each offer (Kacelnik et al., 2011). These two decisions may be implemented by separate, possibly interacting, diffusion-to-bound processes. These ideas are somewhat well-supported by reaction time data (Freidin et al., 2009; Shapiro et al., 2008). Another implication of this idea is that in choice, options are given special status: default (the currently attended one) and alternative (the other one). A good deal of evidence supports the idea that cortical choice processes adopt this framing (Kolling et al., 2012; Boorman et al., 2009; Boorman et al., 2013; Kolling et al., 2014; Azab and Hayden, 2017).

*Implications for the framework:* If we attend single offers in turn and accept or reject each one, direct comparison of values *per se* need not occur (Kacelnik et al., 2011). Comparison may instead result indirectly from the fact that we cannot choose both options if both options are favored for an ‘accept’ decision. A form of value comparison may sneak in via the accept-reject process if the accept-reject decision is made relative to the background, and the background consists of the other option (or best of others in the case of more-than-two-option choice). As a consequence, we do not need separate pools of neurons for representing the two offers, nor do we need additional neurons to perform the comparison **(Figure 1)**. Getting rid of comparator neurons avoids the difficult binding problem by which the offer-selective neurons are dynamically configured to converge on specific comparator neurons. One advantage of using a single pool is that the brain can use all its resources to the difficult problem of value estimation, which requires sensory information, memory and prospection, rather than using anatomically separate computational resources for every option that is (or even that can be) available. The disadvantage, of course, is that the serial nature of value estimation can slow down the decision-making process and requires memory.

**Figure 1.**
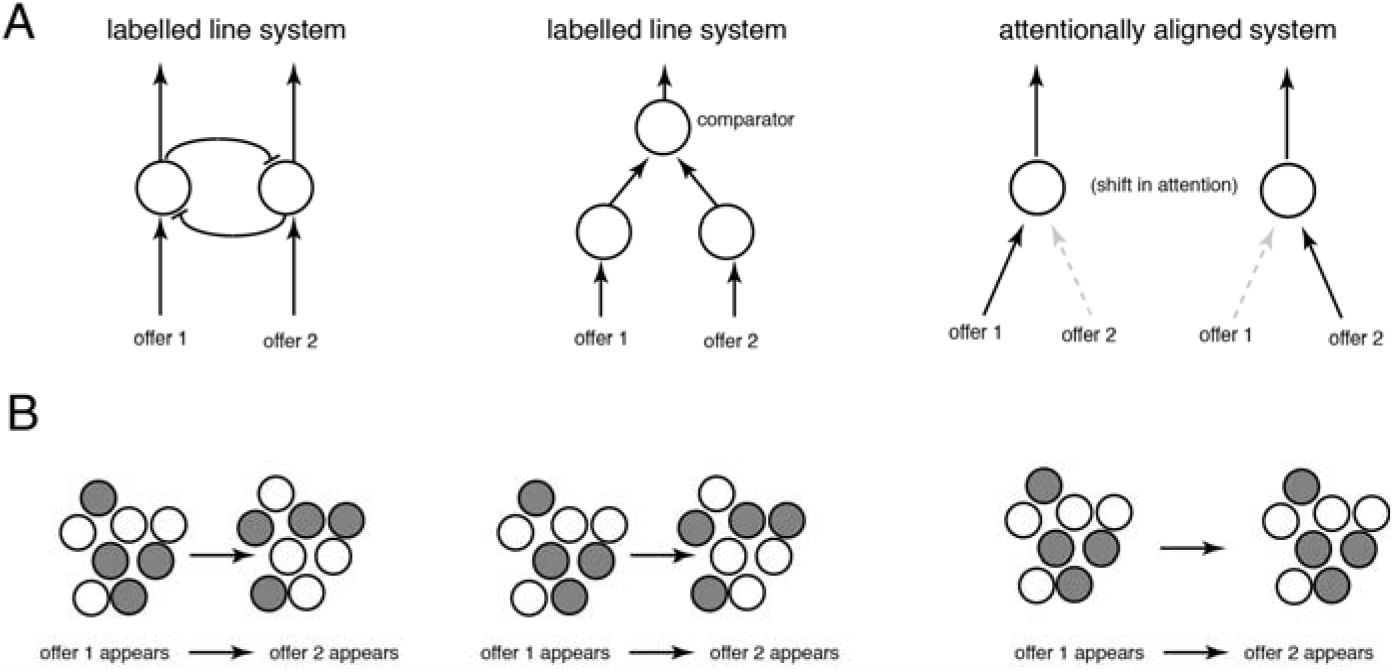
Illustration of core ideas of one- and two-pool models. **A**. *Left:* One approach to modeling choice is the labeled line approach. Each neuron is associated with a specific option and these neurons compete, typically through mutual inhibition, for control of behavior. *Center:* Another take on the labeled line approach has three classes of neurons, one for each option and a third for comparison. *Right:* An alternative *attentionally aligned* approach (supported by recent work reviewed here) eschews labeled lines, but instead involves alternations between states corresponding to just one offer. When attention shifts, the inputs to the value neurons change to reflect the attended option. Because only one option is attended, value sensitive neurons do not need to have information about which option their value signals. **B**. When attention shifts from one option to another, a labeled line system will switch which neurons are strongly activated; an attentionally aligned system does not. Nor will tuning functions change. Thus, measuring which neurons are involved in signaling the values of the two offers, and their tuning, can test between two-pool and one-pool models.

**Figure 2.**
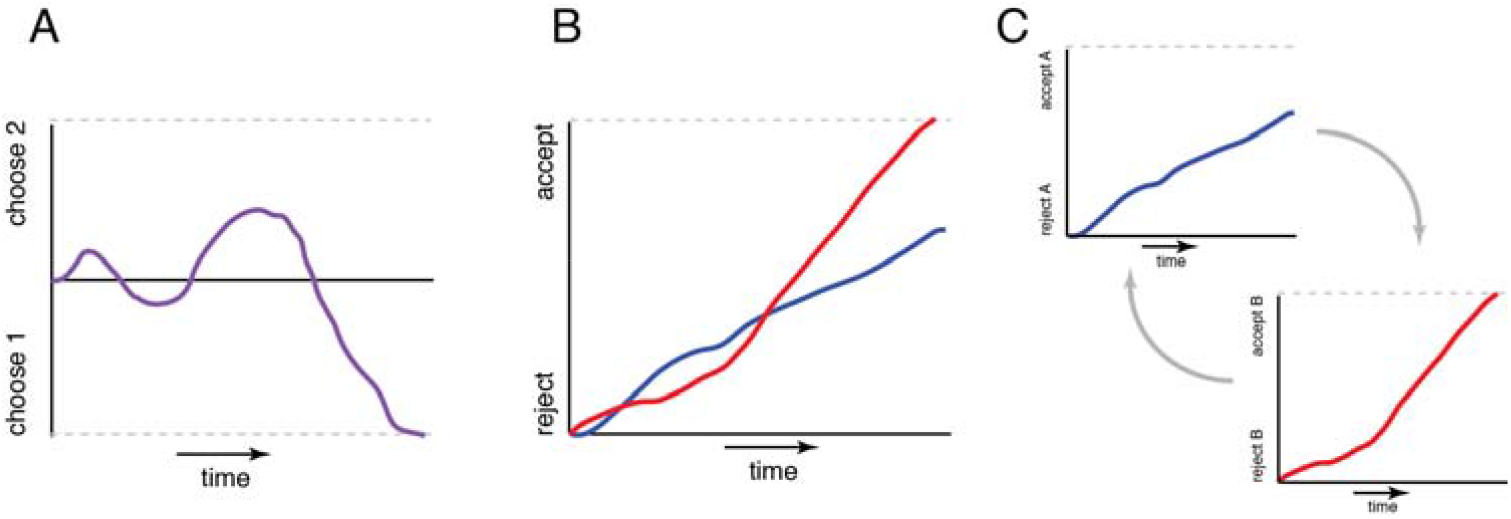
Illustration of basic framework of models. **A**. In simultaneous choice (“tug of war”) there is one decision variable that drifts between two bounds, corresponding to choice of option 1 and 2, respectively. **B**. In independent choice (“sequential choice” or horse race) models there are two decision variables drifting between two potentially similar sets of bounds. They may interact or they may not; generally one threshold hit stops the deliberation process and the second one is halted. **C**. Some recent evidence is consistent with a pair of single diffusion processes, only one of which occurs at a time, as determined by the focus of attention. The threshold in the single diffusion process is likely influenced by the background value (estimate of the value of rejecting), which in turn can be determined by the value of the other option and by the value of further exploration.

### 3. Attention, not labeled lines, determines how value is bound to options

When attention shifts from one option to another, value-coding neurons in several regions switch from encoding the value of the first option to encoding the value of the second. Often, neuronal responses are consistent with use of the same *format* to encode offer values across shifts in attention. That is, a neuron positively tuned for the value of the first considered offer will remain positively tuned for the value of the second and vice versa. We introduce the term *attentionally aligned coding* to refer to this response pattern, which can be distinguished from *labeled line coding*, where neuron will change value polarity depending on the option (Azab and Hayden, 2017). Specifically, the term attentionally aligned means that the referent of a value neuron’s firing rate is not consistently aligned to a single option, but rather is aligned to the value of any option within the focus of attention. Attentionally aligned coding is convenient if attention is limited to a single option at a time, but becomes unwieldy if multiple options can be attended at once.

This pattern was anticipated in neuroimaging studies (Lim et al., 2011), and in careful studies of behavior (Krajbich et al., 2010). An attentional alignment has been reported in neurons in vmPFC, VS, OFC, dACC, and subgenual ACC (sgACC) (Rudebeck et al., 2013; Blanchard et al., 2015; Xie et al, 2017; Rich and Wallis, 2016; Azab and Hayden, 2016; Azab and Hayden, 2017; Strait et al., 2014; Strait et al., 2015), and is consistent with another recent OFC study (McGinty et al., 2016.) Evidence for attentional alignment is illustrated in **Figure 3**. As will be discussed below, attentional alignment is not a novel idea, but rather the same basic principle by which feature coding in the ventral stream works (Desimone and Duncan, 1995).

**Figure 3.**
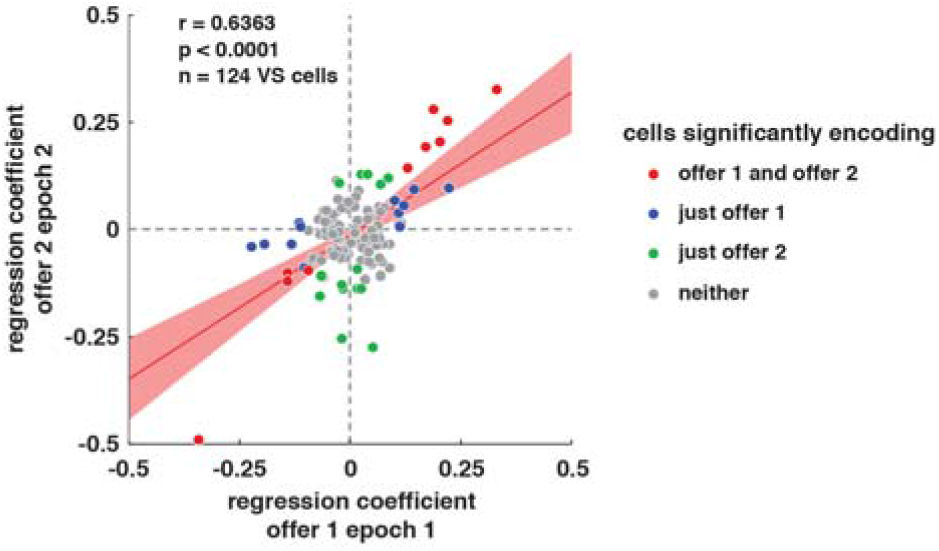
Neurons use a similar ensemble coding format (signed regression coefficient) for the values of offer 1 and offer 2 when they are attended in sequence. In this illustration, options were presented and thus attended asynchronously, and regression coefficients for each neuron were estimated in the appropriate epochs. Each dot indicates a single neuron; its x and y position indicate the linear component of its tuning function for each option. The positive correlation between the two indicates a preservation of tuning, as predicted by a single-population model. A labeled line model would predict that points would cluster around the anti-diagonal and produce an anti-correlation. Data from VS are shown (Strait et al., 2015); similar patterns were observed in other core value regions (see text).

*Implications for the framework:* If value coding is attentionally aligned, then the framework can have a set of value-sensitive units that are ignorant of the details of the input stimuli. Specific value-sensitive units in the network have an organizational advantage: they will not need to be precisely wired with offer layer neurons. This arrangement gives the system much more flexibility to deal with rapidly changing options, new options, and more than two options. One disadvantage is that if an ensemble of attentionally aligned neurons uses the same format to encode the value of two different options, a decoder cannot know, without some additional information (specifically, which option is attended), to which option a neuron firing rate refers. By contrast, in a labeled line coding system, there is no ambiguity about which option a neuron’s firing rate indicates: after all, the line is labeled. On the other hand, if the decoder knows the status of attention, then the referent of the neuron’s firing rate is unambiguous. Thus, the option-value binding problem can be solved without need for any supervisory system other than the one that controls attention.

### 4. One pool of neurons, not two

When attention shifts, and the value code shifts with it, a good deal of evidence indicates (or at least is consistent with the idea) that it is the same neurons activated for the previous option that are activated for the next one (Rudebeck et al., 2013; Azab and Hayden, 2017; Blanchard et al., 2016; Blanchard et al., 2015; Xie et al., 2017; Rich and Wallis, 2016). In other words, the brain may use only one pool of neurons to encode the two different values at different times, not two separate ones. At least one study indicates that some of these regions use a single pool of neurons to encode offered and chosen values as well (Blanchard et al., 2016).

A simple test for separate populations is to compare unsigned regression coefficients (this is similar to, but more statistically sensitive than, performing a Venn Diagram analysis, **Figure 4**). This method reveals evidence in favor of a single population in OFC, vmPFC, VS, and dACC (Azab and Hayden, 2016; Blanchard et al., 2015; Strait et al., 2014; Straight et al., 2015; Wang and Hayden, 2017). A more sensitive method uses Bayesian statistics to ask whether the tuning functions for the two variables supports a single or dual clusters (Blanchard et al., 2016). This method rejects any option-specific clustering 4 brain regions (VS, vmPFC, OFC, and dACC). It also rejects clustering for offer and chosen values.

**Figure 4.**
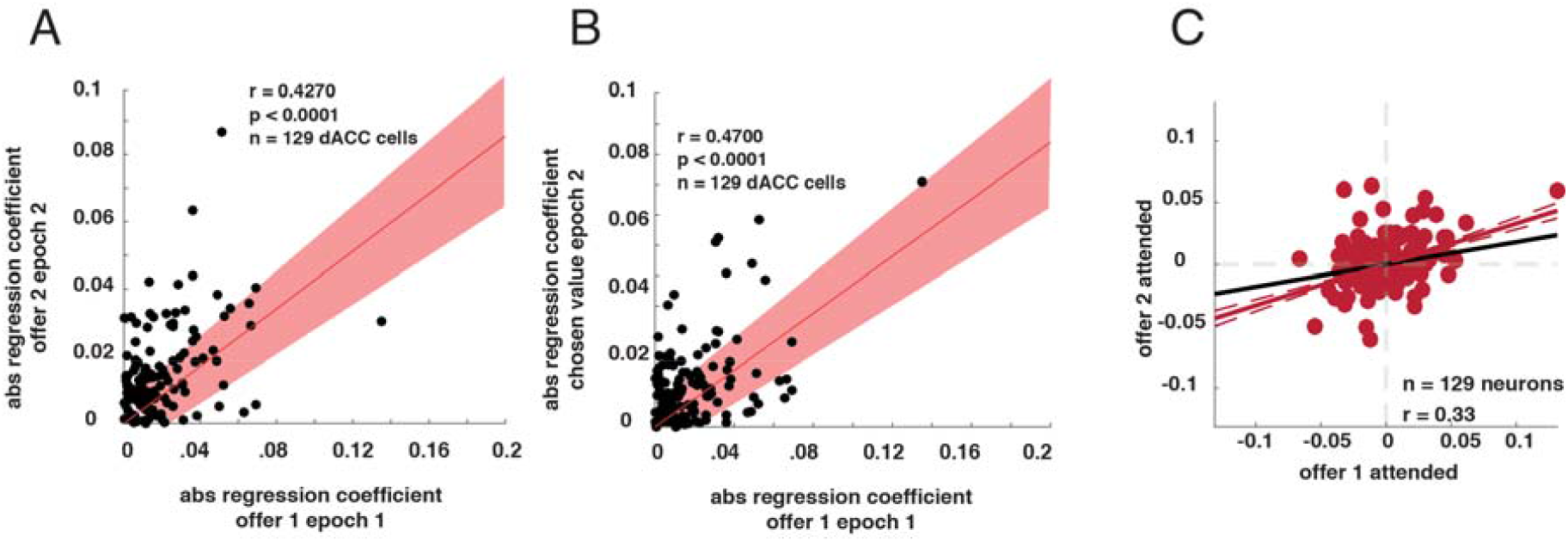
Some evidence for a one-pool model. **A**. In a simple gambling task with two offers presented asynchronously, neurons selective for the value of the first offer are more likely to be selective for the value of the second as well. Selectivity here is measured by the absolute value of the regression coefficient of firing rate against the value of the offer. The positive correlation between the variables indicates that a neuron driven by the value of the first offer is more, not less, likely, to be driven by the value of the second one. We see no evidence of separate populations of cells, as would be predicted by a labeled line model. Illustrative data from dACC shown (Azab and Hayden, 2016); similar patterns were observed in other regions (see text). **B**. Evidence for a single population encoding offered values and chosen values. Illustrative data from dACC shown (Azab and Hayden, 2016); **C**. Aligned encoding of attended offer values. Values of attended offers are encoded in correlated formats across time, in contrast to two-pool model predictions of mutual inhibition through time (Azab and Hayden, 2017).

Note that the case here is not definitive; there is a good deal of ostensibly contradictory empirical support for two pools, and several papers for data consistent with a two pool model (Padoa-Schioppa, 2011) The question of how many pools there are is difficult to answer because the brain may in principle divide up the two offers in any of a number of ways, perhaps arbitrarily and perhaps randomly from trial to trial. Methods that average across multiple trials may then therefore average across the two pools making two look like one. Our analyses so far suggest that neurons do not consistently align to the first/second offer or the left/right offer in asynchronous left-right choices (Azab and Hayden, 2017; Blanchard et al., 2017; Blanchard et al., 2015). Perhaps the strongest evidence so far comes from datasets with simultaneously recorded cells, allowing robust single trial analysis, which still fail to indicate separate pools of cells (Rich and Wallis, 2016).

*Implications for the framework:* The one-pool finding, if true, goes hand in glove with the attentional alignment hypothesis. Specifically, if there is a single pool of neurons, its firing rates must somehow be linked with an option. The most straightforward way would be to limit reference to a single possible option, defined by attention. That system would allow the set of neurons to flexibly encode the value of any offer, and would free the system from having to have a rigid linkage for offers and values. The narrow restriction of attention to a single option would thereby resolve the option-value binding problem. This logic also works for the tentative finding that offered and chosen values are encoded by the same neurons: presumably the chosen offer is attended around and immediately after the time it is chosen, and so it should be encoded in the same neurons that encoded its value at offer time.

### 5. Opposed tuning for offer pairs at the time of choice

At the time the second offer is attended (and the value of the first is available in working memory), the decision-maker can begin comparing their values. At this time, firing rates of neurons in several regions encode the difference in values of the two offers. Specifically, individual neurons tend to show opposing tuning functions for their values. These regions include vmPFC (Strait et al. 14), VS (Strait et al., **Figure 5**), dACC (Azab, 2017), and sgACC (Azab and Hayden 2016), as well as the PMd (Pastor-Bernier Neuron) and SEF (Chen and Stuphorn, 2016). These signals are observed in the same neurons that encode the values of the individual offers and not a separate class of neurons. This pattern is broadly consistent with the finding that several brain regions show coding for the difference in the values of the two offers (Basten 2010; Boorman et al., 2009; Fitzgerald et al., 2009; Hunt et al. 2012). The value difference is *the* key decision variable for economic choice – a simple threshold applied to value difference will produce a choice. It is thus, arguably, a signature of value comparison.

**Figure 5.**
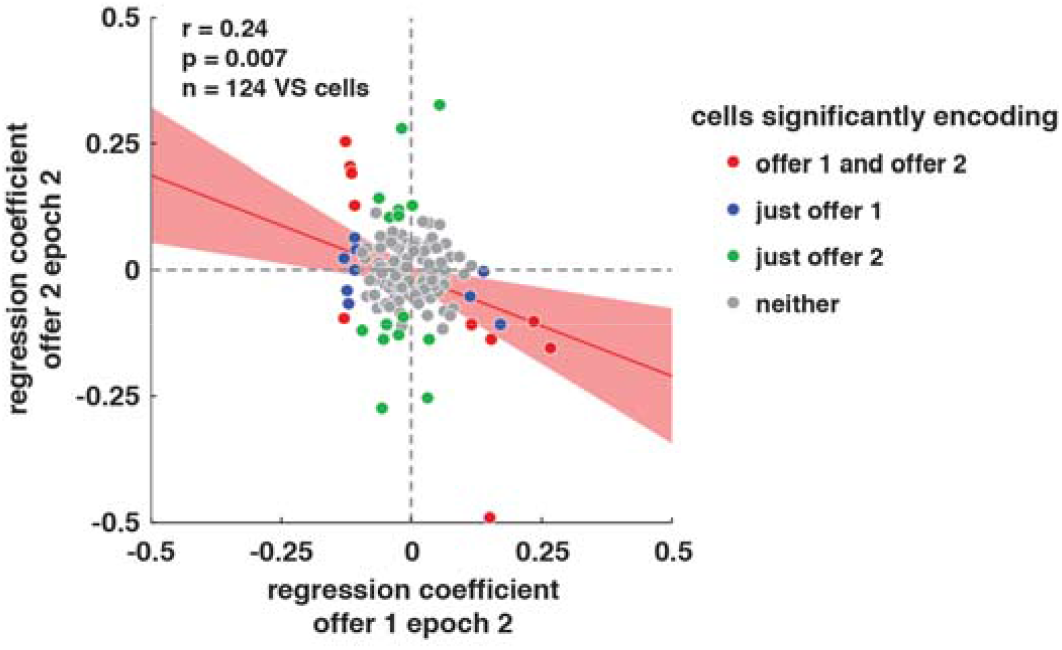
Neurons use inverted tuning formats (reversed regression coefficients) to encode the values of the two offers during choice. Each point indicates a single neuron. The x- and y- positions of the dots indicate the linear term of the regression coefficient for the firing rate of that neuron at the time of choice against the values of the two different (and uncorrelated) offers. The negative correlation indicates that these coefficients are anti-correlated and thus that the population encodes the difference in the values of the two offers. Illustrative data from VS shown (Strait et al., 2015); similar patterns were observed in other core reward regions.

*Implications for the framework:* Putative value comparison signals, in the same neurons that encode values of offers, indicate these neurons do not specialize in encoding the value of the attended offer. Instead, they have a more sophisticated and flexible role in choice. Specifically, they can encode the difference (or a function thereof) between the the attended and the remembered values. Doing so requires them to have some kind of working memory store (whether active or passive) and raises the question of what form it takes (see model below). Note also that the presence of value comparison signals in multiple regions suggests that the comparison occurs simultaneously in multiple regions (see Discussion).

### 6. Activation of motor plans during deliberation

When we evaluate options, and before we choose, the anticipated motor plans of both option are encoded in premotor and parietal cortices (Cisek and Kalaska, 2005; Scherberger and Anderson, 2007; Baumann and Scherberger, 2009; McPeek and Keller, 2002). When the action is clear and overt, that action plan is called an affordance (Cisek, 2007). We use the more generic term *action plan* here to mean pretty much the same thing as affordance, but to include contexts in which the action plan is not clear (imagine for example you are asked which entrée you wish to order, but there is no menu to point at). As evidence accumulates in favor of one option, its corresponding action plan gets stronger and the other one gets weaker, until the decision threshold is reached (Cisek and Kalaska, 2005; Cisek, 2006). The intensity of a given action plan is positively correlated with its value relative to the other one (Pastor-Bernier 2011; Cisek, 2012). The gradual evolution of these processes, in turn, gives rise to decisional commitment (Thura and Cisek, 2014). Together, these findings support a biased competition model for economic choice, which extends classic ideas of biased competition from the perceptual system to the motor system (Cisek, 2005; Cisek and Kalaska, 2010; Pastor-Bernier 2011).

*Implications for the framework:* These results suggest that attending to one offer will activate its action plan and that switching to the other will suppress its action plan and enhance the other’s. During deliberation, these modulations will not trigger an action, but they will be critical for the process of selection and commitment that occur when deliberation ends. These findings also raise the possibility of a solution to the action binding problem as well: if the attended offer activates its corresponding representation in the premotor system, then there is no ambiguity about which option that action code corresponds to.

## PART II. A proposed framework for the neural basis of economic choice

The recent empirical findings provide important new constraints for any proposed mechanisms for the neural basis of choice. These constraints, in turn, point towards the following general framework, which accounts for the major threads of data described above. The goal of this review is to sketch out this framework. We do not pretend the framework is complete. It will need to be developed and revised with new, more detailed exploration and data.

We assume that the brain’s decision-making system can be thought of in the following manner **(Figure 6)**. Sensory inputs activate specific populations of units that represent complex features, and thus the activation of those features in the so-called offer layer defines the identity and characteristics of the offer under the scrutiny of attention. Thus, the offer layer has a distributed representation of offer in the visual field, including their locations. Each unit in the offer layer corresponds to one or more (in the likely case of mixed selectivity) features of possible options. These offer-layer units then activate a representation of the option’s value in units in a separate value layer. Responses from the value layer convey no information about the identity of the option; they simply signal the value of the currently attended option. (Note that we use the term value for convenience. The variable could be any variable or set of variables that correlates with choosing the attended option, such as “evidence in favor,” or signals that reflect the values of individual object features).

**Figure 6.**
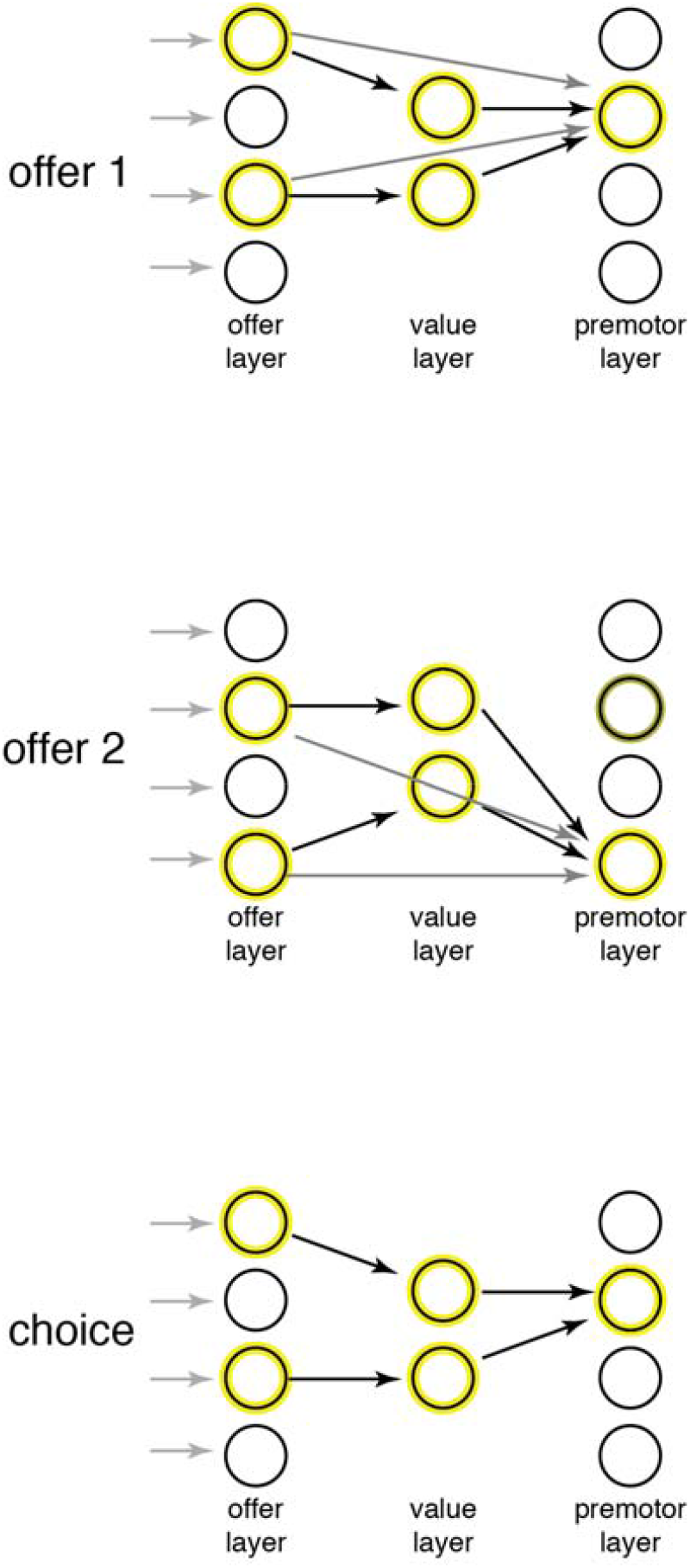
Our proposed framework in practice. When the **first offer** appears, its feature detectors are excited, which define the distributed response in the offer layer. These activate the appropriate value units in the appropriate manner to signal the value of the first offer. They also activate the corresponding premotor layer units (gray arrows). Those are the units that, if the action they signal is released, the animal will choose the offer. When the **second offer** appears, its feature detectors are excited. They active the appropriate value units, which are likely the same ones that were activated by the first offer, and with the same tuning function. They also activate their appropriate premotor units. Finally, **following choice**, the chosen offer is attended and so its features, value, and action units are activated. This activation allows for credit assignment processes to know the appropriate elements to sculpt for learning.

The activation of the offer layer will also activate the option’s action plan in a premotor layer. The action plan is the specific action that would be used to select the option, and can be as specific as a reach or as abstract as the concept of “select this option using the appropriate action when that action is later identified.” The premotor units get signals from both the offer and the value layer. The option information activates the associated action; the value information activates all output actions non-specifically (arrows are not displayed but they are understood as present), providing a general drive to act. The interaction between the value units and the offer units allows for only the attended action to be selected. Finally, the framework allows for additional non-specific modulatory inputs to all action units, which lets extraneous factors such as urgency to affect the likelihood that an action will be triggered (not shown).

Within the context of a choice task, units in the value layer take on the property of *response-dependent suppression* (although this property can in principle be in other layers of the circuit, meaning its response to the first attended offer attenuates its response to the second one in proportion to its response to the first (or a function thereof, more generally). Suppression is not necessary per se, as response-dependent enhancement or any other time of temporal dependence of past responses could certainly work as well. However, response-dependent suppression is a prominent feature of the inferotemporal cortex (IT, e.g. Miller 1991) and may be observed in the reward system as well (e.g. Barron et al., 2013) and dependence on previous outcomes is commonly observed in the reward system (e.g. Kennerley et al., 2011; Hayden et al., 2011), reasons for which we favor that neuronal implementation of our framework. This response-dependent suppression will serve the purpose of a within-cell memory (i.e. doesn’t require an additional external memory buffer and thus can occur within one pool) that will produce value comparisons. There are many possible neuronal mechanisms that could implement response-dependent suppression; we use a simple one for concreteness, one for which there is ample evidence in different domains and thus can be a computational motive: divisive normalization (Carandini and Heeger, 2012; Reynolds and Heeger, 2009).

Note that the idea of repetition suppression, regardless of its specific neuronal implementation, makes a specific novel prediction in asynchronous choice contexts: that regression weights for the second offer computed in the second epoch will be reduced relative to regression weights for the first offer computed in the first epoch. To test this prediction, we performed a new analysis on the relevant datasets we have collected (Strait et al., 2014; Strait et al, 2015; Azab and Hayden, 2016; Blanchard et al., 2015a). Consistent with our prediction, we found that the average unsigned regression weight for the second offer in the second epoch was lower than the weight for the first offer in the first epoch (vmPFC: β1=0.0049, β2=0.0029; VS: β1=0.016, β2=0.010; OFC: β1=0.040, β2=0.021; dACC: β1=0.02, β2=0.0139; sgACC: β1=0.0136, β2=0.0121). These differences are statistically significant in all cases (p<0.001 for vmPFC, OFC, and dACC; p=0.004 for VS; p=0.012 for sgACC). Thus, although the assumptions of the model would need to be revised in the future, at least there is some compelling evidence in its support and concedes further exploration of its implications.

### First fixation: value and action plan for first option

Given this general description of the framework, we next walk through the steps of choice. We propose that in practice, consideration of options is nearly always asynchronous. That is, even when multiple options are presented simultaneously, attention selects one of them for scrutiny first, possibly covertly (Krajbich et al., 2010). When the first offer is attended, the units responsive to its features and/or identity in the offer (feature) layer are activated; these units proceed to activate corresponding value and action units. The action will not yet be triggered. In most cases, assuming the need to decide is not extremely urgent, the first option is likely to be automatically rejected in order to consider the second option; this would be implemented by the global modulatory inputs. Specifically, we can say it would be rejected because the background benefit-cost ratio is quite high: it includes the informational value of the second offer at the very low time and energetic costs of simply looking at it.

### Second fixation: relative value and action plan for second option

When attention shifts to the second offer, its corresponding offer-layer units will be activated. These units then activate the corresponding value-layer units – which will be the same ones that signaled the value of the first offer; they will also use the same format to do so (e.g. a unit with positive tuning for offer 1 will have positive tuning for offer 2). However, when the second offer is processed, the value-layer units will continue to show response-dependent suppression for the value of the first offer. If the first offer was particularly good, the response to even a good second offer will be attenuated. If the first offer was poor, the response to the second will be less attenuated. The value-coding units will therefore exhibit simultaneous and anti-correlated tuning for the values of both offers (as in Strait et al., 2014). Notably, the value of the second offer will not be encoded *per se*. It will only ever be encoded relative to the value of the first. The second offer will also lead directly to the activation of its action plan, just as the first offer did. The action plan for offer 2 will be more strongly activated than the one for offer 1, because attention enhances the action plan. However, we anticipate the action plan for offer 1 will be moderately activated, due to system hysteresis. Both action plans will therefore be activated simultaneously (as in Cisek et al., 2005).

Subsequent to the second fixation, subjects may select it or they may return to the first offer. A return to the first offer will lead the value-layer units to encode its value relative to the value of the second. (This hypothesis has not yet been tested, but follows naturally from our framework). Its action plan will be also be enhanced. This process can continue back and forth until an option is selected (as in Rich and Wallis, 2016). Why would a decision-maker come back a second time rather than just decide immediately? We hypothesize that additional bouts of consideration provide a more accurate estimate of the value of the offers to due accumulated response-suppression of the value unit, allowing for fine discrimination of closely valued options.

### Choice and outcome periods: relative value and action plan of chosen option

An option is selected when the activation in the action layer crosses some threshold. The threshold is determined by several factors that combine to determine the value of rejection. There is very little data on the process of threshold computation (but see Kolling et al., 2014). However, we assume that rejection has a high value following the first offer (because of the informational benefit and low cost of inspecting the second one.) The value of rejection will decrease as time increases and the opportunity cost of further deliberation rises. Once one action plan crosses a threshold, commitment occurs and the selected action inhibits other activated actions, so as to ensure only one action plan is implemented (Thura and Cisek, 14.) The selection process leads the chosen option into the focus of attention. As such, its offer units are preferentially activated and value-layer units encode its value. Note that there are no chosen value units; the units encoding the chosen value are the same value-selective units that were involved in choice.

### After selection

After selection, the reward is received, the chosen option will be attended, and its corresponding offer, value, and action units will be correspondingly activated (or reactivated). In addition, post-reward processes will come into play. These post-reward processes include monitoring, learning, adjustment, and updating of priors, as well as possibly switching to new strategies or rules. These processes are unique to the post-reward period, and will therefore create patterns that are not observed in the offer period, but that will be superimposed on the standard offer-related signals (Wang and Hayden, 2017; Nogueira et al, Nat Comm, 2017).

### Extending the framework to more than two options

Our framework deals well with binary choices, but they need an additional feature to handle choices with more than two options (which we call multi-option choices for convenience). Our model here will be more speculative since we do not have unit data from multi-option choices, although relevant lesion (Noonan et al., 2010), neuroimaging (Boorman et al., 2011; Boorman et al., 2013) and perceptual decision-making (Churchland et al., 2008) data exist. We propose that when attention falls on the third option, the brain encodes its value relative to the value of the best of the first two options (Boorman et al., 2013). The brain could maintain a separate buffer to store the value of the best-so-far option, but we propose a simpler alternative with a single unlabeled value buffer.

Specifically, we propose that the brain maintains an active salience buffer – a representation of the entire option space (both the visual scene and some abstract set of options could be included). The buffer tracks the location of the most valued option so far – but not its value, nor its identity or action plan. The computational framework described above can also be extended to account for multi-option choices using the idea of the salience buffer. The basic idea is that only the offer with the highest value so far is actively remembered, causing divisive normalization on the current stimulus being attended (cf. Louie et al., 2011). Finally, based on the current response, a choice needs to be made between the new stimulus and the past stimulus with the highest value. This pairwise comparison can be made in the same way as described in the previous section.

### A possible neuronal implementation

The key element of our computational model is the presence of a memory mechanism that affects the response of the value layer to the second option and depends on the value of the first option. We propose that this computation is implemented by repetition suppression, via divisive normalization (Carandini and Heeger, 2012; Reynolds and Heeger, 2009). Note that repetition suppression *per se* is not necessary; similar effects can be obtained if neurons exhibit repetition enhancement. We focus here on repetition suppression because it is strongly supported empirically (e.g., firing rate adaptation).

We assume that the attended option encoded in the offer layer delivers information to the value-encoding layer (see Figure 7). In response to the first offer, with value *V_t_*, the firing rate of neuron *i* (*i* = 1…*N*) in the value layer is

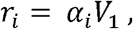

where *α_i_* is a positive coefficient. We consider for simplicity only positive values, although this framework can be naturally extended to negative values too, and to any arbitrary form of tuning curves (e.g., non-linear).

**Figure 7.**
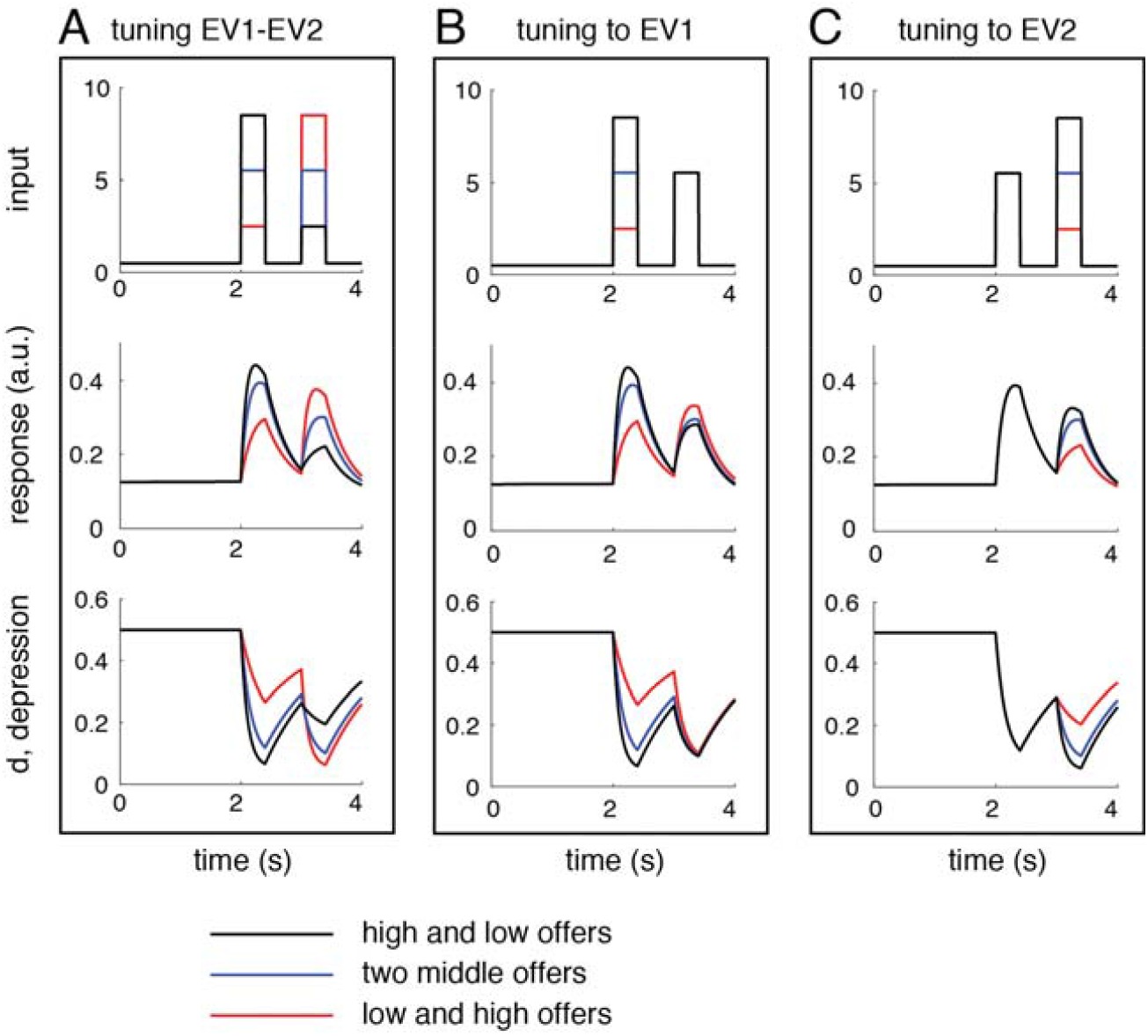
A neuron in the value layer has similar tuning to the values of the first and second offers, and shows repetition suppression. **A**. The tuning to the difference between offer values (tuning to EV1-EV2). **B**. to the first value (tuning to EV1) and **C**. the second value (tuning to EV2) are shown. In B and C, the values of the first or second offers are fixed, respectively. The input (top row), the response (middle row) and the depression variable (bottom row) are displayed as a function of time. The input, encoded in the projections from the offer to the value layers, is proportional to the expected values of the two offers, presented at times 2 sec and 3 sec. respectively. Three different conditions are used (black, blue and gray lines), see text.

When the second offer appears, the response of the value-encoding neurons will be diminished because of repetition suppression in the value (or even in the offer) layer. We assume here that repetition suppression in the value layer is caused by divisive normalization of the neuronal response to the second option in proportion to the strength of the first response. Specifically, the response of neuron *i* in the value layer to the second offer becomes

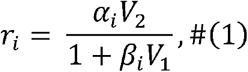

where *β_i_* is a small positive number (0 < *β*_1_ ≪ 1). As a consequence, responses to the second offer will be reduced even if the values of the two options are identical, consistent with the experimental data provided above.

Finally, a choice between the first and second options needs to be made based on the responses that are available in the final stage, that is, the responses to the second offer in Eq. (1). The choice cannot be done with a homogeneous layer of value-encoding neurons, but it can be done with a heterogeneous layer where the sensitivity parameters *α_i_* and *β_i_* differ across neurons. This is because setting a threshold to Eq. (1) with identical parameter values for all neurons causes biases in the choice by preferring the first offer over the second one, or vice versa. However, this bias can be avoided by simply linearly combining the responses of a heterogeneous population of *N* neurons to make it approximately equal to the difference of values of the first and second stimuli:

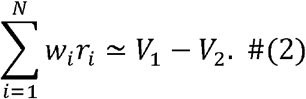

This linear combination can deliver a very good approximation of the actual difference if neurons are heterogeneous and the population is sufficiently large, as we will see in a neuronal implementation of this basic algorithm (see **Figure 8**). This is because if the divisive normalization parameter *β_i_* in Eq. (1) is small, the firing rates can be expanded approximately as linear function of both values. Combining several of those approximately linear functions, it is possible to compute the value difference, which is again a linear function. Therefore, a layer of value neurons with repetition suppression has all necessary information to perform a sequential comparison of two offers, and this information can be extracted by a simple linear readout.

**Figure 8.**
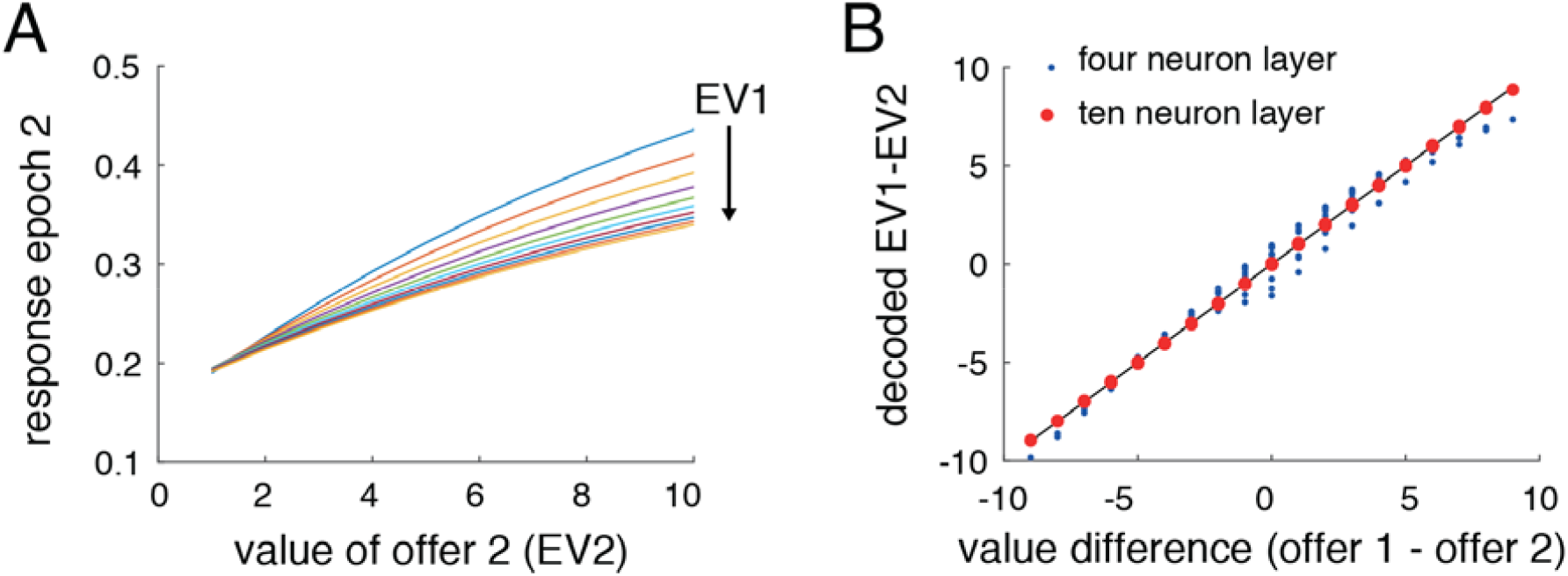
A heterogeneous population of neurons in the value layer can faithfully encode the value difference between first and second offers. **A**. The response of a representative neuron in the value layer during the second epoch increases with the value of the offer in that epoch, EV2, but it is also negatively modulated by the value of the offer previously presented, EV1. This mixed encoding makes impossible to read the value difference from just a single neuron. **B**. Decoded value difference as a function of the real value difference for a population of four (blue) and ten (red) neurons. The decoded values get closer to the actual values (unit slope line, black) as the population is larger. The decoder is based on a linear readout of the population using the responses at 100 ms after second offer onset, trained using linear regression, Eq. (2).

What kind of signals in the brain could carry information from the past to the present in a format that allows also comparing values of sequentially attended stimuli? One such potential candidate is synaptic depression (Tsodyks and Markram, 1997; Abbott, 1997). Synaptic depression acts on the inputs to a neuronal population in such a way that continuous stimulation causes synaptic resources, such as number of vesicles and amount of neurotransmitter, to be depleted. Due to its slow decay, depressing synapses can hold information in working memory for several seconds (Mongillo et al., 2008; Miller and Desimone, 1994; Miller et al., 1991). Thus, synaptic depression is a potential mechanism for facilitating the comparison between the values of two offers presented asynchronously through repetition suppression. It is possible that there are multiple mechanisms with similar effects working together, for instance, firing rate adaptation in the offer and value layers. Here we show simply that synaptic depression is a good starting point, although we acknowledge that, due to its rigidity in the slow time scales involved, it will insufficient to accommodate the large variations of timing in which decision making can occur. This proposal then should be seen as a workable example of how these changes may occur, illustrating the viability of our framework.

We consider a value layer comprised of *N* independent neurons described by their temporally modulated averaged firing rate, *r_i_*(*t*) (*i* = 1…*N*) receiving inputs subject to synaptic depression. Each neuron in the value layer receives the external input *I_i_*(*t*), modeled as a time-varying signal weighted by the value of the stimulus plus background activity, *I*(*t*) = *a_i_* + *b_i_* × *V*(*t*). We assume that the value of the attended stimulus *V*(*t*) is computed as a linear combination of activities in the offer layer (see **Figure 6**), although of course non-linear function beyond linear can be achieved by multi-layer networks. The net input into each neuron is computed as *d_i_*(*t*) × *I_i_*(*t*), where *d_i_*(*t*) is the synaptic depression variable for the inputs for neuron *i*, with takes lower values the higher the activity was in the recent past. Therefore, the net input into each neuron is not simply the external current, but a normalized version of it with a normalization factor that depends on the previous history of attended options and their values. Further details for the models are described in the Methods section. The dynamics of a neuron in the value layer is shown in **Figure 7** for three relevant scenarios. The external input to the neuron is shown in the top panel, while the response and the synaptic depression variables are shown in the middle and bottom panels respectively.

In the first scenario (tuning to EV1-EV2; **Figure 7A**), the external input alternates between a high and a low value offers (black, top panel), two intermediate value offers (blue) and one low and a high value offers (red). The response of the neurons follows the same trend as the input (middle panel), while the synaptic depression variables displays the reversed trend (lower panel). Note the response to the intermediate values (blue, middle panel) in the first and second epochs: the response is reduced during the second stimulation epoch compared to that in the first epoch, even though the stimulus value is identical in the two conditions. This phenomenon corresponds to repetition suppression, as implemented by divisive normalization (Methods, Eq. 4).

This can be understood by looking at the temporal evolution of synaptic depression variable. Initially, this variable has a relatively large value due to low spontaneous firing rate (around one half). However, during attention to the first option, the external input increases and as a consequence the depression variable decays to a lower value (blue, lower panel). Right after offer offset, the depression variable starts to recover and increase towards the basal value. However, the increase is slow due to the long recovery time constant of synaptic resources, and thus the depression variable does not have time to reach the basal value. Indeed, when the second offer is attended, the depression variable has a value that is lower than the basal value. This difference leads to a response to the second offer that depends on the value of the first offer.

In the second scenario (tuning to EV1, **Figure 7B**), the value of the first offer ranges from high (black, top panel) to intermediate (blue) and low (red), while the value of the second attended offer is fixed at an intermediate value. During the presentation of the first offer, the response of the neuron increases with its value (middle panel), indicating that this neuron has a positive encoding of the first offer value. What it is interesting to observe is that during the presentation of the second offer, this cell is still tuned to the value of the offer attended in the first place. However, the tuning is reversed: higher responses are obtained for the lower value in the first attended offer. Also the tuning to the value of the first attended option during the second epoch is reduced compared to the tuning during the first epoch. These two patterns reproduce the experimental results from our lab in vmPFC, VS, and dACC (Strait et al, 2014; Strait et al., 2015; Azab and Hayden, 2016) and echo those of Pastor-Bernier and Cisek (2011).

In the final scenario (tuning to EV2), the value of the first offer is fixed, while the value of the second offer varies. Consistent with experimental results (Strait et al, 2014; Strait et al., 2015; Azab and Hayden, 2016), the tuning of this cell to the value of the second offer is positive. Thus, the neuron tends to keep the same polarity towards stimulus value regardless of stimulus identity or presentation timing.

### Decoding choices

We next asked whether a downstream decoder can make an accurate choice based on the activity of the neurons in the value layer during the second offer epoch. As noted, the response to the second attended option is inverted relative to the first. This inverted tuning allows the system to compare the values. How can this information be extracted? As with the computational framework, it is not possible to read out this information if only one type of neuron is present in the value layer. This is because the firing rate of a neuron during the second epoch depends on the values of both first and second attended offers and does not necessarily compute a value difference between the two. Our strategy is then to create a heterogeneous population of neurons in the value layer, which is a realistic feature throughout the brain architecture. Heterogeneity can be introduced by choosing neuron and depression parameters randomly (Methods).

With a value layer consisting of just four neurons, it is possible to estimate the value difference approximately (**Figure 8**, blue points; max error = 0.97). With ten neurons, it is possible to estimate value difference with high precision (**Figure 8**, red points; max error = 0.08). Although these simulations are based on deterministic dynamics, the presence of response variability can be partially alleviated simply by using larger populations if differential correlations are weak (Moreno-Bote et al., 2014). Therefore, it is possible to compare values of sequentially presented offers by linearly reading out the activity of a small neuronal population in the value layer during attention to the second offer.

## DISCUSSION

Here we review recent discoveries about the neuronal correlates of economic choices and propose a novel guiding framework for future models of how that choice occurs. In this framework, only one option is attended at a time and processing of that option leads to either acceptance or rejection. Rejection often leads to consideration (and sometimes choice) of the next option. During deliberation, attention to an option activates a representation of its value and of the action plan associated with choosing it. This action plan can be specific or it can be abstract, that is, it can in principle be a commitment to a proposition (Shadlen and Kiani, 2013). Our framework requires a single pool of value-sensitive units whose responses encode the value of the attended option relative to the value of previously attended options. It does not involve two pools or more of cells that use labeled-line coding and that compete for control of action. Comparison is accomplished through a value normalization process that can occur simultaneously at multiple levels and that may involve a response-dependent suppression of future responding. As such the evaluation, comparison and the selection are made by the same pool of neurons.

The proposal is not meant to be a formal model for choice, but is, rather, to be a general framework that can guide the development of such models in the future. One particular limitation of the framework is that it does not correspond to the unit level. For example, the strict division into an offer layer, a value layer, and an action layer is not supported by current data. Instead, individual cells are likely to have multiple contributions in multiple layers simultaneously. These functions may even change and adjust with task context (e.g. Hunt et al., 2013). Another example is that value-sensitive neurons, such as those in orbitofrontal cortex may be stimulus-specific, and thus not directly analogous to our value layer (e.g. Schoenbaum et al, 1998). A third example of a limitation of our framework is that value comparison is likely to occur not within a single region, but rather through a distributed consensus process that includes ostensibly motor and association regions both (Cisek, 2012; Chen and Stuphorn, 2016; Hunt and Hayden, 2017). Ultimately, we propose that our framework may be a description of the algorithmic level, but not the implementation level, of choice.

### Relation to models of sensory memory-guided decisions

Our framework is partially inspired by well-known models of memory-guided perceptual decisions (Miller, et al., 1991; Miller and Desimone, 1993; Lui and Pasternak, 2011; Hayden and Gallant, 2013; Mirabella et al., 2007; Machens et al., 2005; Romo et al., 2002; Romo and Salinas, 2001; Romo and Salinas, 2003). Typically in memory-guided perceptual decisions, a memorandum is presented to the subject followed by a delay and then a probe. The subject is then asked to perform a perceptual discrimination on the relationship between the memorandum and the probe (e.g., do they match? Which has higher frequency?). One approach to modeling these decisions is to allow the memorandum to modify the sensory tuning properties of neurons so that their response to the probe makes the correct classification automatically (Miller et al., 1991, Machens et al., 2005). This general approach has been successful in modeling mid/high-level and prefrontal responses (with visual memoranda) and prefrontal responses (with somatosensory memoranda.) Indeed, we propose that binary economic and mnemonic decisions may function through similar brain mechanisms.

The overlap between our proposed framework and the framework used for perceptual decisions is not limited to its relationship with memory-guided decisions. The attentionally aligned coding scheme we propose is shared with perceptual systems. For example, neurons in the ventral visual system have large receptive fields that often contain multiple stimuli competing for attention. Focusing attention on a particular stimulus causes the neurons to behave as if the attended option were the only one present. Thus, the identity of the attended option is identified only by the status of attention. When attention shifts (within the receptive field), the tuning stays the same but the response changes to one that is based on the newly attended stimulus.

This principle, known as *biased competition*, has proven successful at explaining responses of neurons in the ventral visual stream and offers a basis for theorizing about memory, attention, and learning (Desimone and Duncan, 1995; Moran and Desimone, 1985; Chelazzi et al., 1998). Our framework is, in essence, an extension of these ideas past the temporal pole, along the uncinate fasciculus, and into the orbital and medial regions of the prefrontal cortex. We are not the first to make this analogy. From the motor side, Cisek and colleagues have argued that biased competition principles also apply to representation and choice signals in motor and premotor regions (Cisek 2007; Cisek and Kalaska, 2010; Pastor-Bernier and Cisek, 2011). We agree with this idea and propose that it extends backwards. One appealing feature of this idea is that it allows the brain to make use of a single principle to make both perceptual and economic decisions, rather that use a completely different architecture for the two types of choices.

### The neuroeconomic binding problems

One virtue of our framework is that it offers a solution to three important neuroeconomic binding problems that are difficult to avoid with labeled-line models. They concern how values are bound to options, to actions, and to choices (Akaishi and Hayden, 2016; Strait et al., 2016; Hare et al., 2011; Cai and Padoa-Schioppa, 2014.) Values must be linked to their corresponding options (the *value binding* problem, Akaishi and Hayden, 2016; Strait et al., 2016). Then, to select that option, we need to link the result of a comparison with the action that will be used to select it (the *action binding* problem, Hare et al., 2011; Strait et al., 2016; Cai and Padoa-Schioppa, 2014). Finally, once the choice is resolved, we need to link the experienced value with the choice that produced it (the *outcome binding* problem). This is one example of the broader class known as credit assignment problems (Sutton and Barto, 1998; Schultz, 2006).

These binding problems can be understood by analogy to the feature binding problem (Treisman and Gelade, 1980; Shadlen and Movshon, 1999; Engel and Singer, 2001). Imagine seeing a red square and a blue circle; how does your brain know that it is not seeing a red circle and a blue square? Neuronal activity encoding each option dimension must be somehow coordinated. This coordination is unlikely to come through specialized neurons that are sensitive to any combination of multiple features – this would lead to a problem of a combinatorial explosion (Von der Malsburg, 1981; Shadlen and Movshon, 1999; Plaut and McClelland, 2010). One possibility is that this problem is solved by the degeneracy introduced by attention: if only one option is attended at a time, then the dimensions can be assumed to be related to the same single object. The same principle may apply for value as well.

These binding problems are difficult to solve in a labeled-line system. If line labels are stable, our brains would need neurons for all possible options; this is unrealistic. If a new option is added to the mix, new neurons would have to be added. Would they be kept in reserve just in case a new option appears? What if ten new options appear at once? This approach would require complex and specific wiring, ready to go for any possible choice. If the labels are assigned dynamically, then in situations with dozens of choices – such as when choosing cereals at the grocery store – we would need competition between dozens of neuron types. This approach would also require an as yet unidentified supervisory system to assign labels and implement the assignment. More importantly, it would not solve the binding problem: how would the decoders know which options had been assigned to which neurons? How could they know which action to perform to select the option? The costs of such coordination are daunting.

In our model, option identity and value / action / choice can be decoded by the principle of degeneracy: if there are only one option, value, and action within the focus of attention, then they can be assumed to correspond. Thus, in our framework, binding is implemented by attention, and is determined solely by temporal context, not by stable labeled lines. By doing so, the potential combinatorial explosion is contained (Von Der Malsburg, 1981; Shadlen and Movshon, 1999). Thus, in our view, the strict bottleneck imposed by limited attentional capacity is a feature, not a bug.

### Unification of stopping and economic choice literature

The neuroscience of economic choice has tended, by and large, to borrow ideas from the neuroscience of perceptual decision-making (e.g. Rorie et al., 2010; Pais et al., 2013; Krajbich et al., 2010; Louie et al., 2011; Hunt et al., 2015). By contrast, it has tended to bypass the equally important literature on stopping and inhibition (Hampshire and Sharpe, 2015; Schall, 2001; Aron et al., 2004). This literature deals with the question of how we plan and prepare actions and how the brain’s supervisory system regulates their expression.

A stopping decision has some conceptual similarities with an accept-reject decision. Both involve a decision about whether to pursue a single action plan or to refrain from it. Neither involves direct comparison of action plans. In a serial choice model, each option is either pursued (accepted) or ignored (rejected). For this reason, serial choice can be likened to as a pair of accept-reject decisions (Kacelnik et al., 2011). We could equally call each accept-rejection decision a stopping decision: accept is go and reject is stop (or withhold from accepting the option). This change in terminology would raise the possibility that economic decisions are, at least in some cases, implemented in fundamentally the same way as stopping decisions.

The benefit of this interpretation is that we already know a great deal about the neuroscience, the pharmacology, the psychology, and even the psychiatry of stopping. It would be extremely powerful if we could import these ideas wholesale into the field of neuroeconomics, and thus gain a good deal of insight in one fell swoop. Thus, for example, the roles of several cortical and subcortical structures are relatively well-known in stopping; if we could predict and test their corresponding roles in economic choice, that would lead to rapid advance in our understanding of choice. For example, the dACC is part of both the canonical economic and the canonical stopping circuitry (Heilbronner and Hayden, 2016). Are these two roles entirely distinct, or are they aspects of a single function? If we can conceptually unify economics and stopping, then we can more adequately answer this question and others like it.

## METHODS

### Description of the neuronal model

The dynamics of the firing rate and synaptic depression variables follow, respectively, the equations

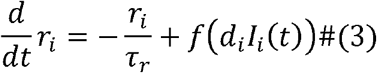

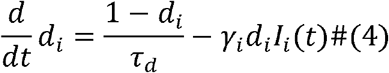

where *f(x)* is the rectified linear function (that is, *f(x) = x* if *x* > 0 and *f(x)* = 0 if *x* ≤ 0) (Tsodyks and Markram, 1997; Abbott et al., 1997). The time constants for the firing rate and synaptic depression dynamics are chosen to be long, *τ_r_* = 500*ms* and *τ_d_* = 2*s*, to allow keeping a memory of the previous stimulus over the interval between stimuli presentation (Mongillo et al., 2008). Long firing rate effective time constants can also be obtained through recurrent dynamics. The firing rate of the neuron in Eq. (3) tracks the total input current with the time constant *τ_r_*. The synaptic depression variable in Eq. (4) depresses whenever the synapse is strongly stimulated, and recovers to its maximum value of one with time constant *τ_d_* if there is no stimulation. How fast the synapse depresses depends on the value of the parameter *γ_i_*.

Heterogeneity of the neuronal populations used in Figure 8 was generated by selecting at random from a normal distribution the tuning parameters *a_i_, b_i_* and the depression parameter (means: [0.5, 1, 1]; standard deviations: [0.05 0.1 0.1], respectively).

## Acknowledgements

This work was supported by an R01 from NIH to BYH (DA037229) and grants PSI2013-44811-P and FLAGERA-PCIN-2015-162-C02-02 from MINECO (Spain) to RMB.

